## Supplementary files for "Diversity within the adenovirus fiber knob hypervariable loops influences primary receptor interactions"

Baker *et al*.

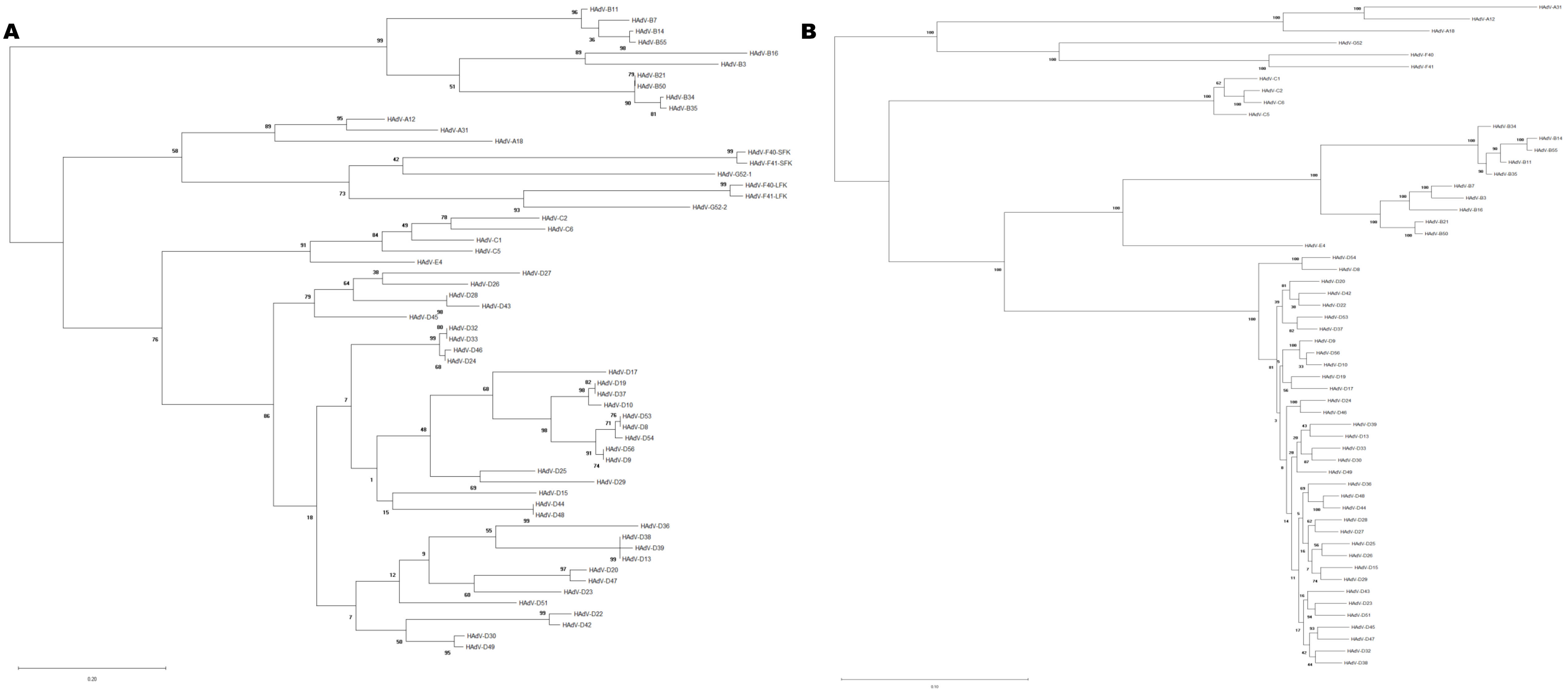


**Supplementary Figure 1: Phylogenetic analysis of adenoviruses by whole genome and fiber-knob domain.** Dendrograms showing uncondensed maximum likelihood trees (percentage confidence shown by numbers next to nodes) were generated from alignments of fiber-knob domain amino acid sequences of adenoviruses 1-56 (A) or whole genome NT sequences (B). While greater diversity can be seen compared to the condensed tree, many nodes are poorly supported.

**Supplementary Figure 2: HAdV-D26 and HAdV-D48 fiber-knob domains are predicted to form more stable trimers that HAdV-C5 fiber-knob but have similar overall topology.** Interface energy calculations performed using PISA are shown on the bar chart with lower values indicating more stable interfaces (A). Amino acid sequence alignment of the 6 tested adenovirus fiber-knob domains is shown with the β-strand regions (underlined) (B). The arrows indicate the positions of the β-strands within the HAdV-C5K structure originally defined by Xia et al (1995), numbering indicates the amino acid position from the first amino acid of that species' fiber-knob protein. n=3, where each calculation is an independent fiber-knob: CAR interface, error bars indicate mean±SD.

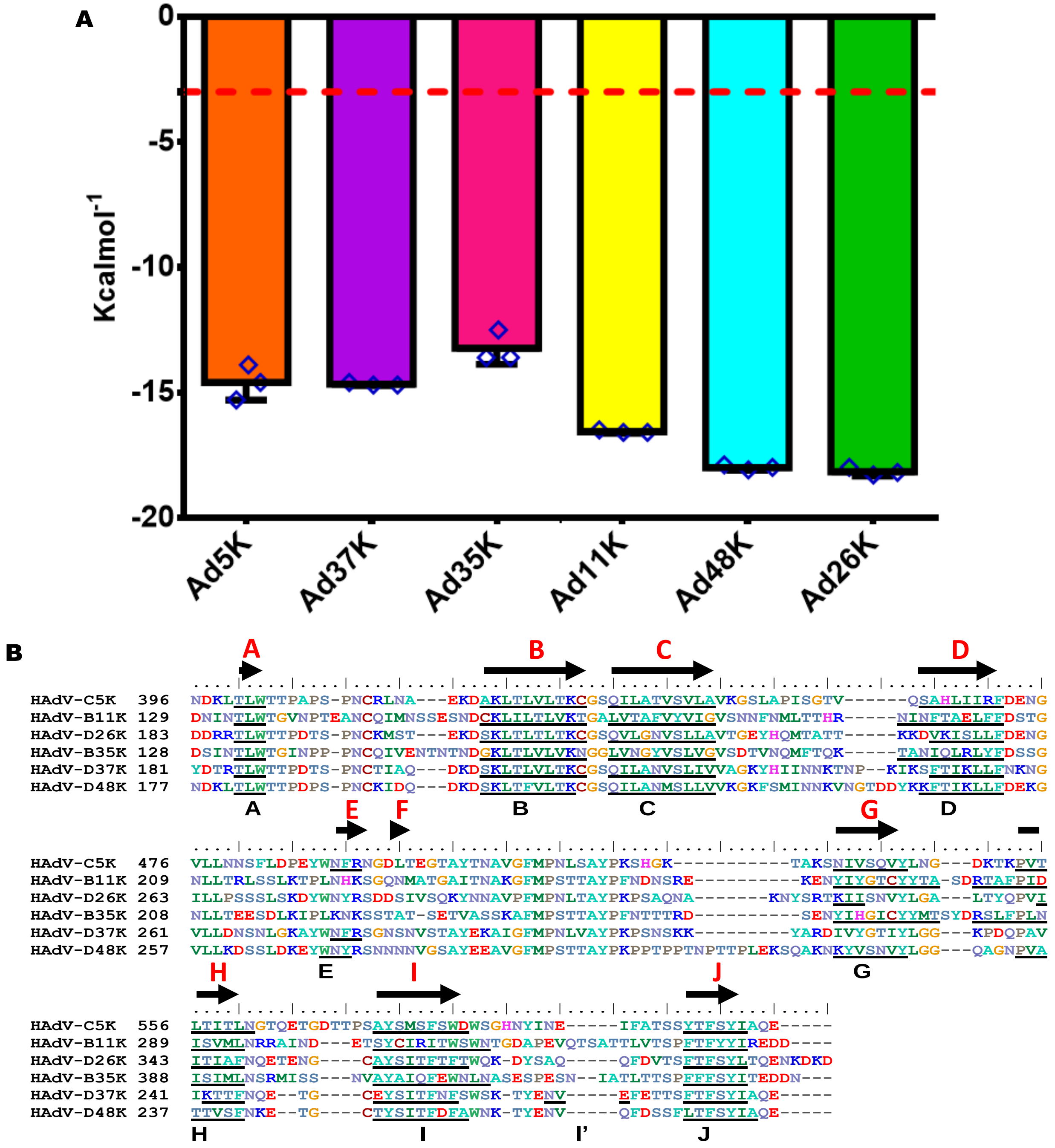

**Supplementary Figure 3: Analysis of polar contacts between HAdV-D26K and HAdV-D48K loops suggests an intricate network supporting specific loop conformations**. The residues comprising the GH and IJ loops of HAdV-D26K (A) and HAdV-D48K (B) are shown diagrammatically. Numbers indicate the position of the loops' terminal residues in the fiber protein. A network of interloop polar interactions is shown by solid lines (one polar bond), and dashed lines (two polar bonds), residues forming part of a helical motif are shaded in blue. The same analysis is seen for the HI, DG, IJ, and CD loops of HAdV-D26K (C) and HAdV-D48K (D).

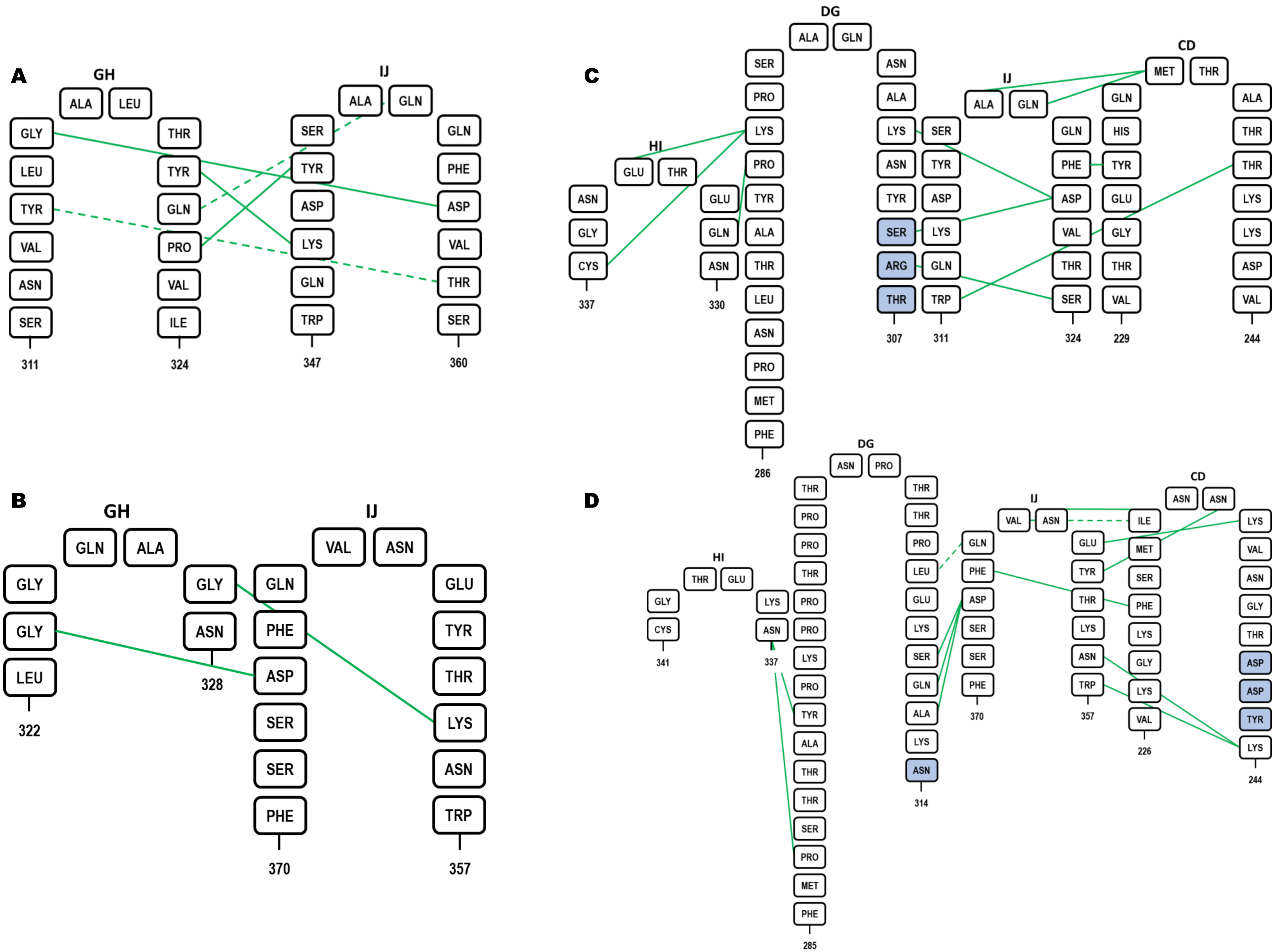

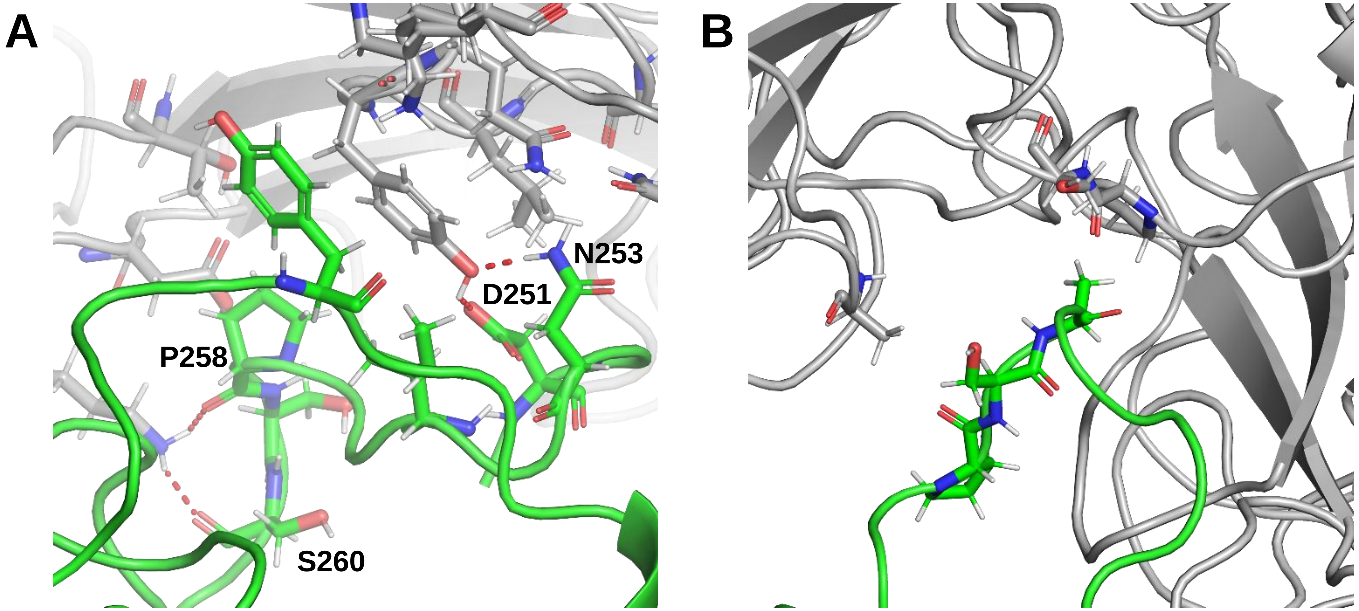


**Supplementary Figure 4: Crystal packing contacts formed by the DG-loop of HAdV-D26K.** Sticks show residues forming crystallographic contacts between the DG-loop of HAdV-D26K (Green) and monomers outside of the biological trimer (Grey). Residues forming polar bonds (Red dashes) are labelled, all other residues are forming Van Der Waals (VDW) contacts at a radius of 3.5Å. Red and blue atoms are oxygen and nitrogen, respectively.

```
HAdV-C5     FMPNLSAYPKSHGK-----------TAKSNIVSQV
HAdV-D45    FMPNLVAYPRPNTPDS-------KIYARSKIVGNV
HAdV-D28    FMPNITAYKP---VNS-------KSYARSHIFGNV
HAdV-D43    FMPNITAYKP---TNS-------KSYARSVIFGNV
HAdV-D26    FMPNLTAYPKPSAQNA-------KNYSRTKIISNV
HAdV-D27    FMPNLVAYPKPTSADA-------KNYSRSKIISNV
HAdV-D25    FMPNLAAYPKSTTTQS-------KLYARNTIFGNI
HAdV-D29    FMPNLLAYAKATTDQS-------KIYARNTIYGNI
HAdV-D17    FMPNLVAYPKPTT-GS-------KKYARDIVYGNI
HAdV-D10    FMPNLVAYPKPSN--S-------KKYARDIVYGTI
HAdV-D19    FMPNLVAYPKPSN--S-------KKYARDIVYGTI
HAdV-D37    FMPNLVAYPKPSN--S-------KKYARDIVYGTI
HAdV-D8     FMPNLVAYPKPTT-GS-------KKYARDIVYGNI
HAdV-D53    FMPNLVAYPKPTT-GS-------KKYARDIVYGNI
HAdV-D54    FMPNLVAYPKPTT-GS-------KKYARDIVYGNI
HAdV-D9     FMPNLVAYPKPTA-GS-------KKYARDIVYGNI
HAdV-D56    FMPNLVAYPKPTA-GS-------KKYARDIVYGNI
HAdV-D13    FMPNLKAYPKPTKTASDK-AENKISSAKNKIVSNF
HAdV-D38    FMPNLKAYPKPTKTASDK-AENKISSAKNKIVSNF
HAdV-D39    FMPNLKAYPKPTKTASDK-AENKVSSAKNKIVSNF
HAdV-D36    FMPSTTAYPKPTNNTSTD-PDKKVSQGKNKIVSNI
HAdV-D51    FMPNLKAYPKNTTTSSTN-PDDKISAGKKNIVSNV
HAdV-D23    FMPNLKAYPNPTTSTTNP-STDKKSNGKNAIVSNV
HAdV-D20    FMPNLKAYPKP--STVLP-STDKNSNGKNTIVSNL
HAdV-D47    FMPNLKAYPNPKTSTVLP-STDKKSNGKNTIVSNL
HAdV-D32    FMPNIKAYPKPTTDTSA-KPEDKKSAAKRYIVSNV
HAdV-D33    FMPNIKAYPKPTTDTSA-KPEDKKSAAKRYIVSNV
HAdV-D24    FMPNIKAYPKPTTDTSA-KPEDKKSAAKRYIVSNV
HAdV-D46    FMPNIKAYPKPSTDTSA-KPEDKKSAAKRYIVSNV
HAdV-D22    FMPNTTAYPKIIDSTTNP--ADKKSSAKKIIVGNV
HAdV-D42    FMPNTTAYPKIINSTTDP--ENKKSSAKKTIVGNV
HAdV-D15    FMPSKTAYPKQTKPT-----NKEISQAKNKIVSNV
HAdV-D44    FMPSTTAYPKPPTPPTNPTTPLEKSQAKNKYVSNV
HAdV-D48    FMPSTTAYPKPPTPPTNPTTPLEKSQAKNKYVSNV
HAdV-D30    FMPNSTAYPKIINNGTAN-PEDKKSAAKKTIVTNV
HAdV-D49    FMPNSTAYPKIINNGTAN-PEDKKSAAKKTIVTNV
```

**Supplementary Figure 5: Species D adenoviruses possess a range of DG loop sequences varying in both length and sequence.** The DG loops of all species D adenoviruses are shown aligned, using clustal omega, to the same region in HAdV-C5 (species C).

```
            ....|....| ....|....| ....|....| ....|....| ....|....| ....|....| ....|....|
HAdV-D37K   TLWTTPDTSP NCTIAQDKDS KLTLVLTKCG SQILANVSLI VVAGKYHIIN NKTNP--KIK SFTIKLLFNK
HAdV-G52SFK TLWTPPTSNP NCTVYTESDS LLSLCLTKCG AHVLGSVSLT GVAGTMTNMA E--------T SLAIEFTFDD
CAV-2       TLWTGPGPSI NGFINDTPVI RCFICLTRDS NLVTVNASFV GE-GGYRIVS PT------QS QFSLIMEFDQ
HAdV-D26K   TLWTTPDTSP NCKMSTEKDS KLTLTLTKCG SQVLGNVSLL AVTGEYHQMT ATT-----KK DVKISLLFDE
HAdV-D48K   TLWTTPDPSP NCKIDQDKDS KLTFVLTKCG SQILANMSLL VVKGKFSMIN NKVNGTDDYK KFTIKLLFDE

            ....|....| ....|....| ....|....| ....|....| ....|....| ....|....| ....|....|
HAdV-D37K   NGVLLDNSNL GK-AYWNFRS GNSN--VSTA YEKAIGFMPN LVAYPKPSNS KK-------- -YARDIVYGT
HAdV-G52SFK TGKLLHSPL- VN-NTFSIRQ GDSP--ASNP TYNALAFMPN STLYARGGSG ---------- -EPRNNYYVQ
CAV-2       FGQLMSTGNI NSTTTWGEKP WGNNTVQPRP SHTWKLCMPN REVYSTPAAT ISR------- ----C-GLDS
HAdV-D26K   NGILLPSSSL SK-DYWNYRS DDSI--VSQK YNNAVPFMPN LTAYPKPSAQ NA-------K NYSRTKIISN
HAdV-D48K   KGVLLKDSSL DK-EYWNYRS NNNN--VGSA YEEAVGFMPS TTAYPKPPTP PTNPTTPLEK SQAKNKYVSN

            ....|....| ....|....| ....|....| ....|....| ....|....| ....|....| ...
HAdV-D37K   IYLGGKPDQP AVIKTTFNQE --TGCEYSIT FNFS-WSKTY ENVEFETTSF TFSYIAQE-- ---
HAdV-G52SFK TYLRGNVQRP ITLTVTFNSA A---TGYSL- -SFK-WTAVV -REKFAAPAT SFCYITEQ-- ---
CAV-2       IAVDGAPSRS IDCMLIINKP K-GVATYTLT FRFLNFNRLS GGTLFKTDVL TFTYVGENQ- ---
HAdV-D26K   VYLGALTYQP VIITIAFNQE TENGCAYSIT FTFT-WQKDY SAQQFDVTSF TFSYLTQENK DKD
HAdV-D48K   VYLGGQAGNP VATTVSFNKE --TGCTYSIT FDFA-WNKTY ENVQFDSSFL TFSYIAQE-- ---
```

**Supplementary Figure 6: Previously described sialic acid using adenovirus fiber-knob sequences aligned to HAdV-D26 and HAdV-D48.** The fiber-knob domains, defined as the conserved TLW hinge motif to the C-terminus, of the known sialic acid utilising HAdV-D37, HAdV-G52 short fiber-knob, and CAV-2, aligned to the fiber-knob domains of HAdV-D26K and HAdV-D48K by clustal omega. Residues highlighted in red have been described as direct contacts between HAdV-D37K and sialic acid. Those in yellow form water bridges to sialic acid, from HAdV-D37K. Those highlighted in blue have been shown to be important to the charge dependent HAdV-G52K to poly-sialic acid interaction, and those in green shown to be involved in CAV-2's interaction with sialic acid.

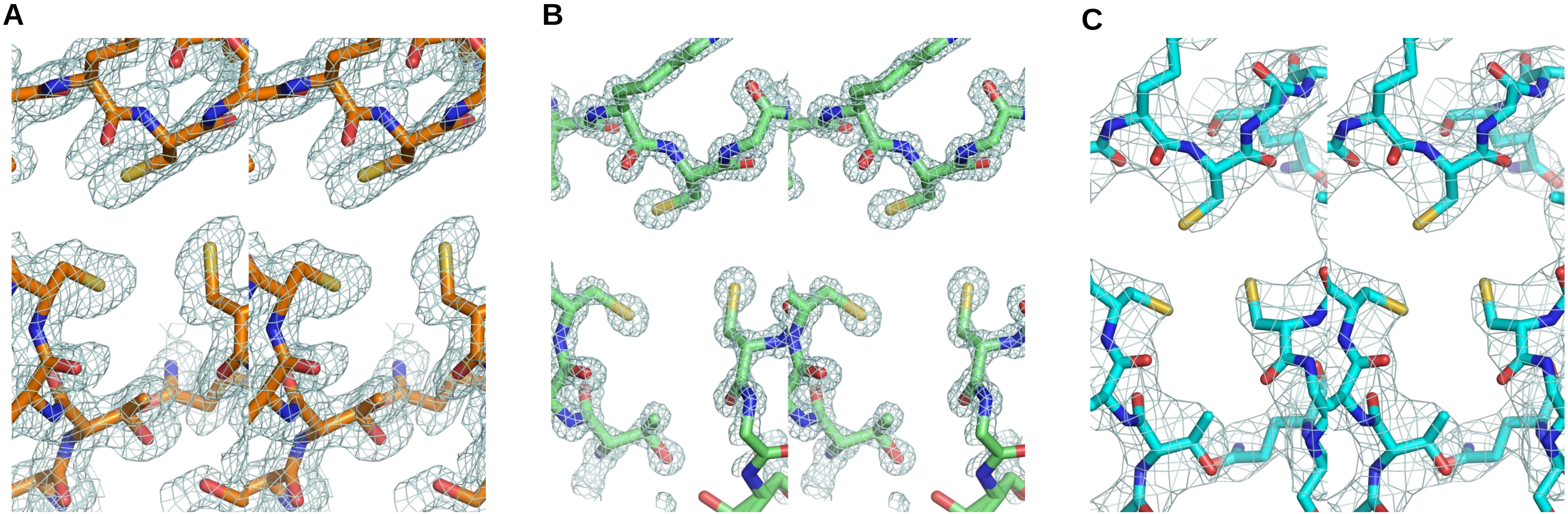


**Supplementary Figure 7: Stereo Images of a representative region of model and map for crystallographic structures determined in this study.** Stereo images of HAdV-C5K (PDB 6HCN) (**A**), HAdV-D26K (PDB 6FJN) (**B**), and HAdV-D48K (PDB 6FJQ) (**C**) are shown coloured in orange, green, and cyan, respectively. Oxygen, nitrogen, and sulphur atoms are coloured in red, blue, and yellow, respectively. The displayed 2Fo-Fc map is contoured at 1.5σ for HAdV-D26K, and at 1.0σ for HAdV-C5K and HAdV-D48K.

**Supplementary Table 1:** Binding energy for each loop of HAdV-D26 and HAdV-D48 resulting from crystal packing contacts, outside of the biological trimer, and the number of polar bonds formed.

| | HAdV-D26K | | HAdV-D48K | |
|---|---|---|---|---|
| **Loop Identity** | **Total Binding Energy (KcalMol$^{-1}$)** | **Number of Polar Contacts** | **Total Binding Energy (KcalMol$^{-1}$)** | **Number of Polar Contacts** |
| **AB** | -0.6 | 0 | N/A | 0 |
| **BC** | N/A | 0 | N/A | 0 |
| **CD** | -0.3 | 2 | -2.6 | 3 |
| **DG** | -6.5 | 4 | -2.4 | 0 |
| **GH** | -3.0 | 2 | N/A | 0 |
| **HI** | N/A | 0 | N/A | 0 |
| **IJ** | N/A | 0 | -0.4 | 0 |

**Supplementary Table 2:** Data collection and refinement statistics for new HAdV-C5, HAdV-D26, and HAdV-D48 Fiber-knob protein crystal structures deposited in the PDB.

| **PDB Entry** | **6FJN** | **6HCN** | **6FJQ** |
|---|---|---|---|
| Diamond Beamline | I04 | I24 | I04 |
| Date | 2017-05-12 | 2018-01-26 | 2017-05-12 |
| Wavelength | 0.9795 | 0.96859 | 0.9795 |
| Crystallisation Conditions | 0.1 M MMT 25% PEGA 1500 | 0.1 M MMT, 25% PEG 1500 | 0.1 M Bis-Tris-propane, 20% PEG 3350, 0.2M NaNO3 |
| pH | 6.0 | 7.0 | 6.5 |
| *a*,*b*,*c* (Å) | 86.01,86.01,86.01 | 102.16,102.44,77.0 | 145.18,145.18,145.18 |
| $\alpha=\beta=\gamma$ (°) | 90.0 | 90.0 | 90.0 |
| Space group | $P\ 2_1\ 3$ | $P\ 2_1\ 2_1\ 2$ | $P\ 4_3\ 3\ 2$ |
| Resolution (Å) | 0.97-60.82 | 1.49-61.56 | 2.91-83.82 |
| Outer shell | 0.97-1.00 | 1.49-1.53 | 2.91-2.99 |
| *R*-merge (%) | 4.3 (74.5) | 13.4 (183.8) | 12.5 (302.6) |
| *R*-meas (%) | 4.5 (97.5) | 15.9 (218.3) | 12.7 (306.3) |
| CC1/2 | 1.00 (0.427) | 0.983 (0.565) | 1.00 (0.705) |
| $I / \sigma(I)$ | 27.3 (0.7) | 7.1 (0.7) | 22.2 (1.7) |
| Completeness (%) | 94.9 (43.9) | 99.8 (99.9) | 100.0 (100.0) |
| Multiplicity | 16.7 (1.6) | 6.6 (6.3) | 41.2 (41.4) |
| Total Measurements | 1,978,768 (6,429) | 876,648 (60,950) | 496,751 (5,136) |
| Unique Reflections | 118,603 (4,055) | 131,951 (9,638) | 12,061 (877) |
| Wilson B-factor($Å^2$) | 8.2 | 18.3 | 74.5 |
| R-work reflections | 112,612 | 125,479 | 11,371 |
| R-free reflections | 5,879 | 6,388 | 572 |
| R-work/R-free (%) | 18.2 / 19.5 | 21.1/23.3 | 20.1 / 29.1 |
| Bond lengths (Å) | 0.025 | 0.011 | 0.019 |
| Bond Angles (°) | 2.339 | 1.534 | 2.293 |
| [1]Coordinate error | 0.020 | 0.087 | 0.370 |
| Mean B value ($Å^2$) | 17.6 | 30.6 | 84.9 |
| Favoured/allowed/ Outliers | 138 / 7 / 1 | 133 / 10 / 1 | 350 / 28 / 7 |
| % | 94.5 / 4.8 / 0.7 | 92.4 / 6.9 / 0.7 | 90.7 / 7.5 / 1.8 |

* One crystal was used for determining each structure.
* Figures in brackets refer to outer resolution shell, where applicable.
[1] Coordinate Estimated Standard Uncertainty in (Å), calculated based on maximum likelihood statistics.

Buffers:

- MMT: DL-Malic acid, MES monohydrate, Tris: pH 4.0-9.0
- SPG: Succinic acid, Sodium phosphate monobasic monohydrate, Glycine: pH 4.0-10.0
